# Resonant Learning in Scale-free Networks

**DOI:** 10.1101/2021.11.10.468065

**Authors:** Samuel Goldman, Maximino Aldana, Philippe Cluzel

## Abstract

Over the last decades, analyses of the connectivity of large biological and artificial networks have identified a common scale-free topology, where few of the network elements, called hubs, control many other network elements. In monitoring the dynamics of networks’ hubs, recent experiments have revealed that they can show behaviors oscillating between ‘on’ and ‘off’ states of activation. Prompted by these observations, we ask whether the existence of oscillatory hubs states could contribute to the emergence of specific network dynamical behaviors. Here, we use Boolean threshold networks with scale-free architecture as representative models to demonstrate how periodic activation of the network hub can provide a network-level advantage in learning specific new dynamical behaviors. First, we find that hub oscillations with distinct periods can induce robust and distinct attractors whose lengths depend upon the hub oscillation period. Second, we determine that a given network can exhibit series of different attractors when we sequentially change the period of hub pulses. Using rounds of evolution and selection, these different attractors could independently learn distinct target functions. We term this network-based learning strategy *resonant learning*, as the emergence of new learned dynamical behaviors depends on the choice of the period of the hub oscillations. Finally, we find that resonant learning leads to convergence towards target behaviors over an order of magnitude faster than standard learning procedures. While it is already known that modular network architecture contributes to learning separate tasks, our results reveal an alternative design principle based on forced oscillations of the network hub.

**SIGNIFICANCE:** Large networks of interconnected components such as genes or machines can coordinate complex behavioral dyamics. One outstanding question has been to identify the design principles that allow networks to learn new behaviors. Here, in simulating learning cycles, we randomly modify the interactions between components and select networks that exhibit a desired behavior. Surprisingly, we find that networks can learn new behaviors faster when the state of the most connected network component is forced to oscillate during learning. Remarkably, using distinct periods of oscillations allows a given network to learn distinct behaviors. While it is known that modular network architecture contributes to learning separate tasks, our results reveal an alternative design principle for which modules are not needed.

## INTRODUCTION

Natural and engineered systems provide a myriad of examples in which large networks of interacting elements must perform well-specified dynamical behavior in a coordinated way (1). While it is often difficult to predict the behavior of such large intricate networks, it is standard practice to use Boolean neural networks to simulate complexity of interactions between network elements (2). However, despite this simplification, Boolean neural networks have emerged as powerful systems that can be trained to learn the dynamical behavior of many technological and biological systems (3). Recurrent connections between network elements, resemble the intricate connectivity with a combination of feedback and feedforward loops observed in biological networks, which can produce a rich array of non-trivial dynamics with fading memory. In contrast with layered, feedforward neural networks, Boolean neural networks with a recurrent topology are well-suited to handle time series data (4).

It has now been well-established that such recurrent networks can be harnessed to mimic complex time-dependent behaviors as diverse as social interactions, gene regulations, electric grids and industrial processes (4, 5). However, despite recent progress, it is still not clear what are the underlying control parameters that govern the flexibility of such networks to learn various behaviors. Here, we aim to identify the conditions that permit a certain class of recurrent Boolean networks to learn multiple target time-series. While we have been initially inspired by recent observations made from experiments with real gene-regulation networks, we limit this work solely to identifying some general properties of idealized networks whose elements are Boolean.

Over the last two decades, a range of experiments have revealed that many large biological networks exhibit a scale-free topology where one network node acts as a hub that governs most of the network activity (6, 7). For example, cellular transcriptional networks have evolved such that a small number of “master regulator” genes can control a large number of downstream genes (8, 9). Recent experimental technologies have enabled several groups to monitor long time series of expression (activation) of these master regulators (10) and to report the existence of pulsatile behavior (11, 12) **(Fig. S1 and S2)**. Overall, these studies suggest that specific timescales of pulses are associated with specific dynamical behaviors of the cell (12-17). Inspired by these experimental observations, we ask how the oscillatory behavior of highly connected nodes (hereafter called *hubs*) such as master regulators, can shape the behavior of large non-linear dynamical systems. Additionally, because it is established that random Boolean threshold networks (RBTN) with scale-free network topologies are able to learn target behaviors more quickly than homogeneous, well-distributed architectures, we focus the rest of this study on recurrent networks with scale-free topologies (18, 19). We demonstrate that distinct dynamical behaviors of RBTN can be pre-selected by adjusting the oscillatory period of the hub node. Furthermore, using an evolutionary algorithm we find that these networks can learn specified target behaviors more efficiently in the presence of an oscillating hub. We term this new property of network-based learning in the presence of input oscillations, *resonant learning*.

Previous work has shown that noise and small perturbations can drastically change the outcome of networks. Stern used Kauffman networks to do demonstrate that “noisy inputs” repeated over several generations of learning rounds could be incorporated by the network to imprint the system a desired behavior (20). From a control perspective, Cornelius et al. demonstrate a strategy to use perturbations to guide a network toward a desired state when only a small set of accessible nodes in the network can be modified, parallel to real world settings such as therapeutic design, when only a subset of genes or gene products can be drugged effectively to “save” the network (21). A separate line of investigations has revealed how oscillatory inputs, often referred to as “frequency-modulated” signals, can be interpreted in small circuit settings. Gao et al. consider how small subnetwork motifs could decode complex temporal signals, and Rue et al. observe how individual nodes in Boolean networks are capable of relaying the frequency of an incoming temporal signal (22, 23). However, to our knowledge, the effect of oscillatory input signals on network learning has not yet been explored. It is not clear either how sustained oscillatory input signals on hub nodes as observed in experiments can shape the dynamics of large dynamical systems such as RBTNs.

### Network Model

We use random Boolean threshold networks (RBTN) as prototypes for the study of large recurrent dynamical systems (4, 24). RBTN’s have been used to mimick gene expression patterns in some organisms (25, 26). We define networks with *N* nodes, {*σ*_1_, *σ* _2_, …, *σ*_*N*_} and a set of directed edges,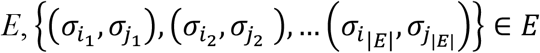. We choose scale-free network topologies with out-degree connectivity distributed according to the power-law distribution, *p*(*k*) ∝ *k*^−*γ*^, which confers the desirable property of “hubs” shared by a large number of biological networks (27-29) (**Fig. 1A**). For example, in regulatory transcriptional networks, these hubs could represent central endogenous regulators such as sigma factors. Moreover, we showed that scale-free networks evolve more efficiently toward a target function than homogeneous networks (Erdös-Renyi), which makes scale-free networks potentially good candidates to identify learning properties of large dynamical systems (18, 19).

**Figure 1.**
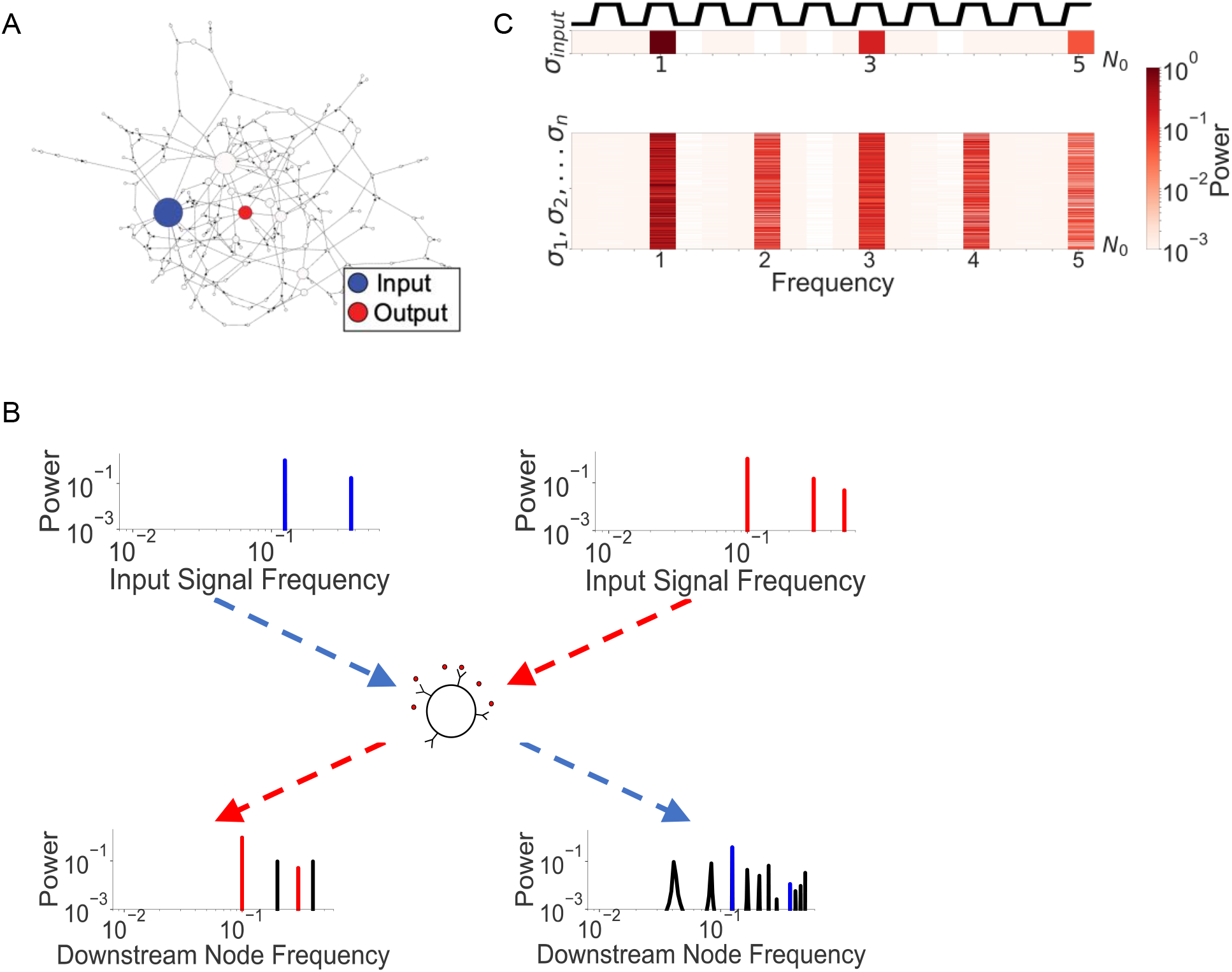
Effect of hub oscillations on downstream nodes. **(A) Illustrative example** of network topology from which we analyze how certain downstream node (red) respond to a given oscillatory behavior of the hub (blue). (**B) Single Node Frequency Response**. Power spectrum (bottom) calculated from a randomly selected downstream node’s time series when hub node oscillates at specific frequencies (top). We note that the plotted frequency here is directly related to the period, 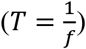. The power spectra of output nodes exhibit both higher and lower frequencies than the input signal. When the input signal has a period of T = 8 (Blue), the network produces harmonics and subharmonics. On the other hand, when the input period is *T=10* (red), the downstream node only exhibits harmonics. We show the corresponding time domain sequences in the SI (**Fig. S5)**. (**C) Input-output relationship network response**. We take this same network and observe the frequency response of all non-frozen nodes (rows) in the network when the input hub oscillates with a frequency of *f = 0*.*1*, equivalent to *T = 10*. We show the input square wave time series and its associated power spectrum above the output power spectrum for clarity. We normalized the frequency axis with the the input frequency of the hub, *N*_0_ = *f* = 0.1, showing that harmonics dominate the behavior of downstream nodes. We repeat this procedure for input period *T = 8* in the SI (**Fig. S6**).

Next, the network dynamics is defined such that the state {0,1} of any node at a given time step correspond to *OFF* and *ON* states of the node respectively. The state of the network at time *t*, which we refer to as ***σ***(***t***), is the state at time *t* of all the genes: ***σ***(***t***) = {*σ*_1_(*t*), *σ*_2_(*t*), ⋯, *σ*_*N*_(*t*)}. The Boolean threshold networks updating rule is based on a simple threshold activation function (**Fig. S3**). We define the updating rule such that the state of any node at the next time step is determined by the product between the weights of incoming connections and the state of the nodes from which they come:

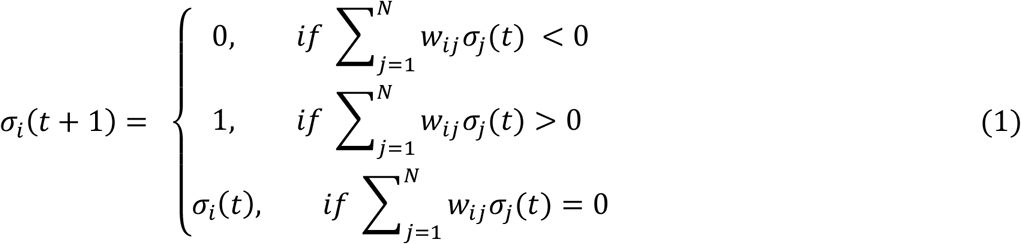

We set the weight *w*_*ij*_ for each directed edge in the network from node *σ*_*i*_ to node *σ*_*j*_ drawn from a uniform distribution on the interval [−1,1], whereas *w*_*ij*_ = 0 if there is no connection between *σ*_*i*_ and *σ*_*j*_. Because the signal that each *σ*_*i*_ receives from its regulators is additive, each weight *w*_*ij*_ exerts either an inhibitory or activating effect on its target node with varying strength. The weights contribute to the updating rule only if the controlling nodes are in the *ON* state. Given the discontinuity apparent in Eq. (1), the dynamical behavior of the network is highly nonlinear. Altogether, this simple non-linear rule is capable of generating a rich variety of dynamical behaviors (25, 26, 30).

## RESULTS

### Hub oscillations propagates in downstream nodes

To probe the effects of hub oscillations on the network nodes and attractor dynamics, we simulate large scale-free networks (*γ* = 1.9) in which the hub node *σ*_*hub*_, the node with highest outgoing connectivity, is forced to oscillate periodically at a pre-set switching rate. we choose *γ* ≈ 1.9 because it corresponds to the critical-like regime of a second order phase transition as shown in **Fig. S4**, where learning algorithms are the most impactful ((19), fig.S10). **Fig. 1A** shows the hub *σ*_*hub*_ in blue, and an arbitrarily node (in red) chosen as the “reporter” or output node. We use periodic and symmetric square waves with period *T* as the driving input signal of the hub (see SI Square Waves and **Fig. 1A**, blue). Surprisingly, we find that distinct time series outputs (attractor cycles) associated with the reporter node (**Fig. 1A**, red) in forced networks can be induced by distinct input periods of oscillation applied to *σ*_*hub*_. Given that *T* only defines the rate at which *σ*_*hub*_ switches between *0* and *1* states, it is surprising that it can induce drastically different network cycles.

To visualize this effect, once the networks has reached an attractor cycle, we monitor the response of single nodes of the network to various input periods *T*. That is, we allow a network to update over many time steps, under the restriction that *σ*_*hub*_ oscillates at a specific period T. Exciting *σ*_*hub*_ at a certain T would yield specific time series for each node *σ*_*i*_ in the network. To evaluate to what degree downstream nodes either transmit the same or different frequencies as those driving the oscillatory hub, we compute the power spectral density (**Fig. S6**) of the time series associated with each node *σ*_*i*_. We can see (**Fig. 1B**) that with an input period *T* = 10, the output node’s time series exhibits the same frequencies as the input node and some other different frequencies.^a^ However, for another input period *T* = 8, the output node responds even with lower frequencies (longer periods) of oscillations than the input node. In general, we find that the period of input oscillations on the hub node can induce a range of downstream behaviors of the output nodes. We summarize this observation using a heatmap that depicts the power spectra for all non-frozen nodes, i.e. those that are switching back and forth between ‘on’ and ‘off’ states) (**Fig. 1C)**. We observe that while the oscillations of the controlling hub have a fixed single timescale, they do not make the downstream nodes of the network oscillate uniformly.

### Hub oscillations induce resonance attractor cycles

After observing that individual nodes respond distinctly to a given input period, we ask whether we can observe more general behaviors when we consider the state of the system at the level of the whole network ***σ***(***t***). Because the updating rule for our networks is deterministic and has finite number of states, the network will eventually enter a fixed cycle of states, which we define as an attractor cycle. We define the start and end of the attractor cycle to be the first two time points at which states, ***σ***(***t***_**1**_) = ***σ***(***t***_**2**_), and all network states prior to time *t*_1_ belong to the transient states of the network (see ***SI Defining an Attractor*** for further specification). We use attractor cycles as means to characterize the overall behavior of the network. We then ask how attractors associated with a given network – the attractor landscape – is perturbed in response to oscillating inputs.

We first specify a null experiment for comparison. Namely, because the oscillating inputs involve switching between the *0* and *1* state of the hub node, we define ground-state attractors as the set of all attractor cycles when the hub node of the network is blocked, made to stay constant, either in the *0* or *1* state. That is, we can sample many different initial conditions for a given network and force *σ*_*hub*_(*t*) = *const*. The network states and the progression they follow over time define the ground-state basins of attraction for a given network. Each network state will eventually converge upon an attractor cycle according to this null reference scheme, and we can quantify how far away each network state is from its corresponding attractor as the number of time steps it takes to converge to an attractor cycle (**Fig. 2A)**. We call this distance the “height” of any given network state. In this scheme, the network states that are part of attractor cycles are considered to have a height of *0* when the hub is blocked to a fixed value. The effect of oscillating the controlling hub becomes analogous to excite the network that bounces back and forth between these ground-state landscapes defined when the network has a hub with a fixed state. Interestingly, we find that this procedure yields new attractor cycles that share some states with the reference basins of attraction but also that have new ones (**Fig. 2A**). As a result, when the hub node is forced to oscillate, the network converges upon new attractor cycles that have non-zero average “height,” as defined above.

**Figure 2.**
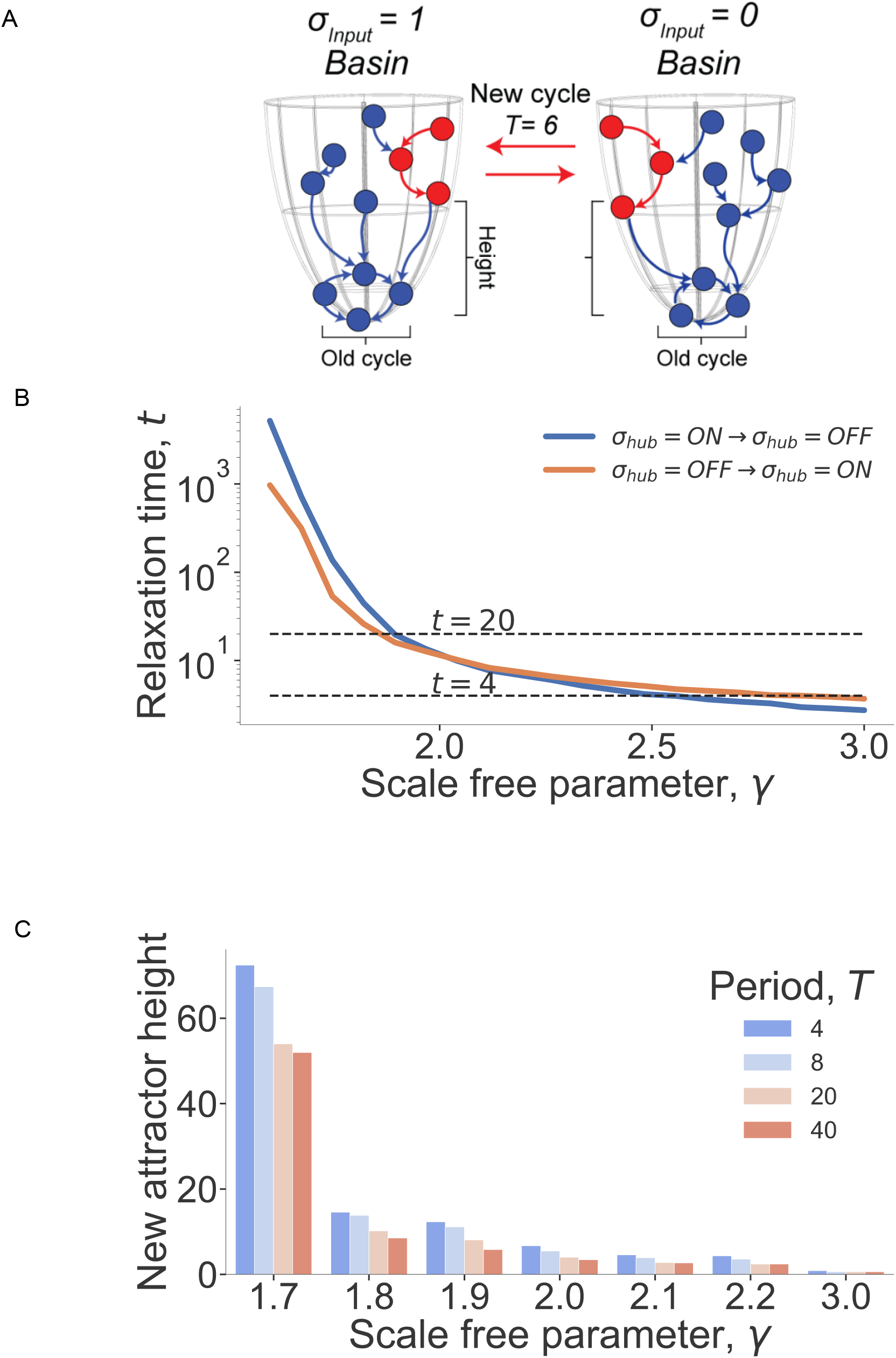
Creating new attractor landscapes. **(A) Oscillating hub creates new attractor cycles**. In a non-oscillating setting, network states (blue) will converge on deterministic attractor cycles (bottom of funnel). However, by oscillating *σ*_*hub*_ we create new attractor cycles (red). New cycles are separated from the ground state cycle by a non-zero height that represents the number of states for the new attractor to relax to the ground state when the hub stops oscillating. (**B) Relaxation time of the network**. We show the average relaxation time, *t*, vs. scale-free parameter *γ* for networks of 1000 nodes. The relaxation time defines the time it takes for the network to settle onto a new ground state cycle when the hub is switched from *σ*_*hub*_ = 1 to 0 (and vice versa). (**C) Heights of new attractors as a function of the scale-free parameter and the period of hub oscillations**. For each T=4, T=8, T=20, and T=40, we average the height of the new attractors from 10 distinct networks, where we probe the landscape with 1,000 random starts. New attractor cycles for more chaotic networks (*γ* = 1.9) yields larger heights i.e. new cycle states are further out in the reference basins.

Next, we characterize the longest oscillation period that can yield new attractor cycles. We reasoned that if the period of switching *σ*_*hub*_between the *0* and *1* states is too long, the network will exhibit the limit behavior with same attractors as if *σ*_*hub*_were either blocked at *0* or *1*; under this condition no new attractor cycle behavior will be reached. To identify the longest possible oscillation period that can yield new attractors, we simulate a random initial condition, block the hub in the *0 (*or *1)* states, and allow the network to fall onto an attractor cycle. Then, we switch the input hub node to the *1* (or *0)* state and measure the relaxation time for the network to reach a new attractor cycle with the new hub state. We repeat this procedure over *1,000* different networks and initial conditions to obtain the average relaxation time for various *γ* exponents. We consider this relaxation time to be an estimate for evaluating the longest period duration beyond which the network will be insensitive to the oscillations of the hub (**Fig. 2B**). Networks with *γ* = 3.0 have relaxation time, t_relax_=4, therefore the longest period for the hub oscillations capable of inducing new behavior is T=2t_relax_=8. Addionnally, we find that the longest relaxation time for networks that have a regime close to critical-like behavior (*γ* = 1.9) to be t_relax_=20. Therefore any periods greater than T=2t_relax_=40 will yield no new attractor cycles and will be associated with a *0* height regardless of how much longer the period is.

Using the average “height” as a metric, we construct a phase-diagram to illustrate how statistically different the new attractor cycles are from the ground-state cycles by generating networks for different values of *γ* in the interval [1.7,3], in the presence of an oscillating *σ*_*hub*_ with different input periods. As expected, the larger the value of *γ* the networks are more ordered and fall on ground-state attractor cycles with height *0*, whereas more chaotic networks with *γ*∼1.9 exhibit more novel attractor cycles composed of network states with greater heights (**Fig. 2C**). Overall, newly generated attractors are in fact novel with respect to each other and with respect to the ground-state attractors as long as the network is sufficiently chaotic with *γ* < 1.9 (**Fig. S7-S8**), which also ensures that different input hub periods can induce different attractor cycles (**Fig. S9)**.

Not only these attractors are selectable by specifying a certain input period, but we find that these newly selected attractors are robust to variations in initial conditions. Specifically, we can estimate this robustness as the number of new attractor cycles identified by sampling a large number of initial conditions. We draw random initial conditions for each network and input period, and we determine the fraction of unique attractors. With *γ* = 1.9 in the chaotic regime, we find that on average very few (fewer than 2%) of the sampled initial conditions give unique attractors (**Fig. S10**). In Boolean scale-free networks, new attractor cycles can be induced by simply changing the duration of square wave periods for a single input hub node. These new attractor cycles are robust to different initial conditions and are mainly governed by the duration of the period of hub oscillations. We call *resonant* attractor cycles, these new network behaviors induced by forcing periodic inputs on the hub.

### Resonant attractor cycles exhibit fast learning dynamics

It has already been well-established that scale-free networks can learn faster than homogeneous networks (19). However, we now ask whether new resonant attractor cycles can also learn target functions. We thereafter a learning scheme similar as (19). We first initialize a population of *N*_*pop*_ = 50 scale-free networks with *γ* = 1.9 (**Fig. S11**). While Oikonomou and Cluzel demonstrated that learning in scale-free networks is not strongly dependent on *γ* (**Fig. S12)** but rather on the general scale-free topology itself, we choose the scale-free parameter *γ* = 1.9 so that networks are near the critical regime (**Fig. S13)** with sufficiently large basins of attraction while maintaining attractor cycles of relatively few states.

The goal of this learning algorithm is to train the network in such a way that a single output node of the network exhibits the same time series of “on” and “off” states as that of the target function, which is a predefined random sequence of 0’s and 1’s of length *L*_*c*_ (**Fig. S14**). Both the output node, which is selected randomly, and the target function remain the same throughout the learning cycles. We perform each simulation for *G* = 10^5^ generations. In each learning generation, we generate *M* = 3 mutated networks from each of the *N*_*pop*_ networks. We score how well all networks, including the unmutated networks, have learned all the target functions. We select the top scoring *N*_*pop*_ networks from the temporary population of size 4*N*_*pop*_ (parent and mutant networks, **Fig. S15**), and repeat this process for all the *G* = 10^5^ generations.

To mutate each network, we use a fixed mutation rate of *μ* = 0.02, such that with probability *μ* we mutate each node in the network, using the same optimized conditions as in (19) with equal probability, a mutation for some node involves changing the weight or target node of an outgoing edge. By only modifying the target node or weight, we guarantee that we maintain the power-law out-degree distribution of the network.

To score the performance of a given network in a generation, we randomly choose an initial condition for the network. Then, oscillating the input node with input period *T*_*c*_, we allow the network to update through time until it falls onto an attractor of length *L*. We take the time series of the output node in this attractor cycle and estimate the distance between this time series versus the target time series of length *L*_*c*_(**Fig. 3**A). Because we want this scoring function to be invariant to the offset of the attractor cycle time series, we consider all circular permutations over the target function. Additionally, it is often the case that *L* ≠ *L*_*c*_, so we repeat both time series to a length *L*′ = *min*(*L, L*_*c*_). By doing this, we avoid penalizing the attractor cycle for a length mismatch with the target function. Thus, we define our score for a single target function of length *L*_*c*_, *Fitness*, as an averaged hamming distance between the target function and the output node’s time series (**Fig. S14**). At each generation, only the *N*_*pop*_ networks (out of 4*N*_*pop*_) with the lowest values of *Fitness* will pass to the next generation **(Fig. S15)**. Furthermore, target functions that are too long are also difficult to learn, especially when their lengths exceed that of the relaxation time of the network. Therefore, we restrict our learning algorithms to input periods, *T < 20* (**Figs. 2B and S16)**. Moreover, we find that the hub oscilation period determines the length of the attractor cycle and that networks learn a target function best when the input period is equal to the length of the target function (**Fig. S17**).

**Figure 3.**
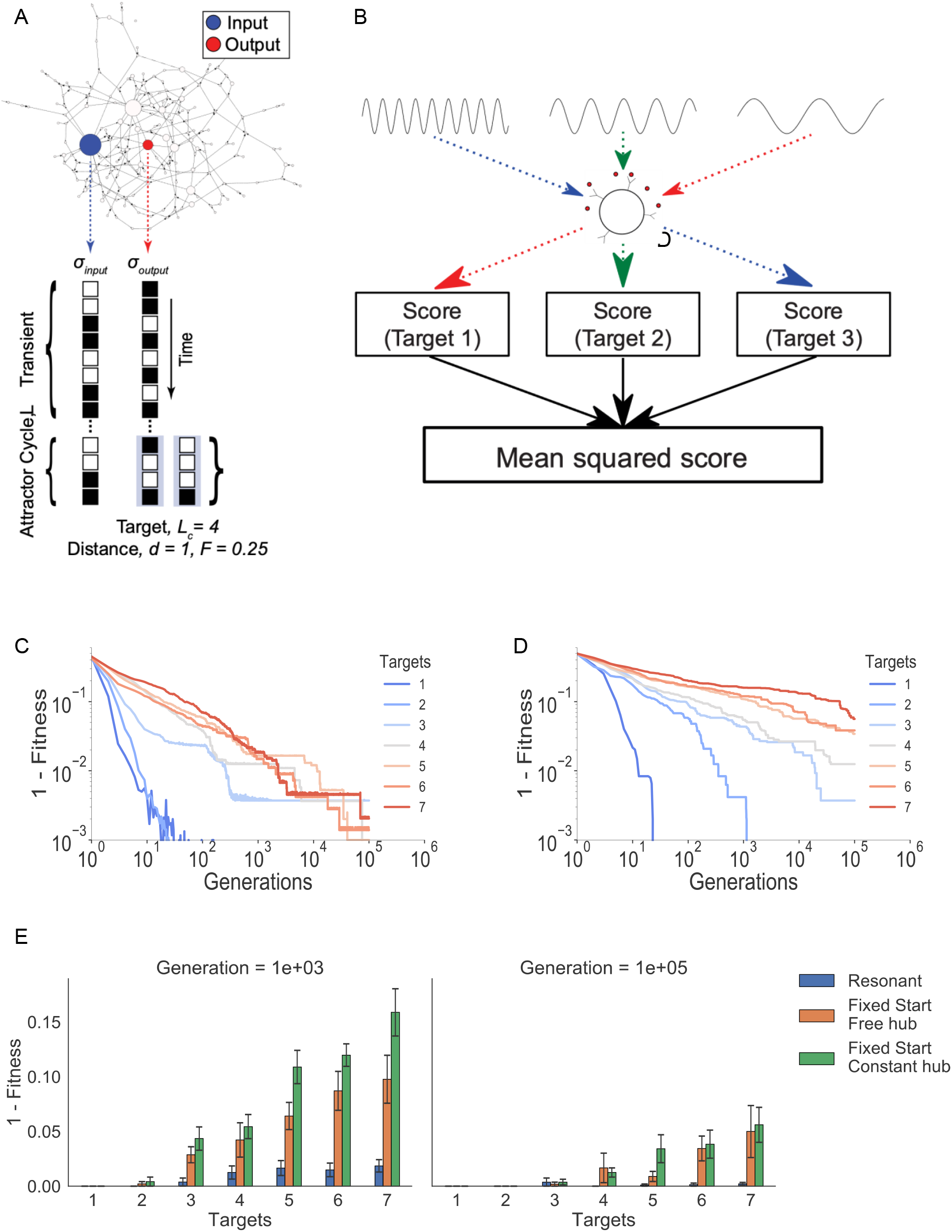
Training networks in the presence of oscillating hub. **(A) Fitness function for learning multiple targets**. We arbitrarily select an output node downstream in the network. For a given initial condition, we score how close the state of this node in the attractor cycle is to a randomly generated target function (see SI for more details**). (B) Learning multiple functions**. We learn multiple random target functions to match various input frequencies. Separately, we run simulations from random initial conditions during which the hub node oscillates successively at all pre-selected input periods, *T*_1_, *T*_2_,. ., *T*_*n*_. In each learning cycle, we score how close the attractor cycle at the output node matches the corresponding target function *f*_1_(*t*), *f*_2_(*t*),. ., *f*_*n*_(*t*). We take the mean squared fitness function across these trials to summarize of how well a network learned multiple behaviors at the output node. **(C) Scale-free networks with oscillating hub can learn multiple target functions**. We run 10 different simulations with *N* = 500, *γ* = 1.9, to show that the network can learn a maximum of 7 different target functions associated with 7 input frequencies before a loss in performance. To decrease variance emerging from trial-to-trial variability, we average over 30 trials for 1-3 targets, 20 trials for 4-5 targets, and 10 trials for 6-7 targets. **(D) Networks with a hub that has a fixed state learn much slower**. Rather than allowing the network to learn in the presence of an oscillating hub with pre-selected period, we provide one fixed state for the hub and one target cycle, similar to the previous learning procedure (forcing the hub node to stay constant). Interestingly, while we might expect the network to learn faster when it must only be concerned with as few as 3 of the 2^500^ total input states, the network learns far slower under this alternative scheme, than in the presence of an oscillatory hub. **(E) Summary plots of results from (C) and (D) for the learning cycle 10**^**3**^ **and 10**^**5**^. We can see that networks with resonant learning scheme are more fit than networks that must learn with a non-oscillating hub, regardless of whether or not the hub node is fixed or allowed to update freely. Bars are plotted with standard error.

Since networks can exhibit different attractor cycles in response to different input hub frequencies, we ask whether these networks are also capable of time-division multiplexing, i.e. learning several target functions corresponding to different input periods. Here, we restrict ourselves to setting the length of the target cycle to be the same as the hub oscillation period when trying to force networks to learn multiple behavior patterns at once. We generate a set of *m* target functions of length *L*_1_, *L*_2_, … *L*_*m*_. In addition, we chose a set of oscillatory input functions with pre-defined periods that will correspond to the length of each target function: *T*_1_, *T*_2_, … *T*_*m*_. Each oscillatory function is applied to the hub node of each network (i.e. the node with highest connectivity). Additionally, to enforce learning multiple functions simultaneously, we take the mean squared error across the different input periods-target functions pairings (**Fig. 3B)**.

We find that these networks can learn as many as seven distinct target functions by simply varying the period of the oscillating hub (**Figs. 3C and S18**). We find that this limit of 7 different target functions is caused by the difficulty for the network to learn both long and short timescale functions simultaneously (**Fig. S16, S18**). The network learns the shorter target lengths faster than the longer ones, but due to the mean squared error scoring mechanism, the network does not unlearn any of its shorter targets to learn the longer targets. We called hereafter this learning procedure *resonant learning*.

We now compare learning efficacy under alternative evolutionary schemes: First, we force the hub node to stay constant, (or to follow the rule in Eq.(1)) but allow the hub node’s outgoing weights to be mutated during learning. Again, we set the simulation to learn up to seven different input initial conditions to match different target lengths (**Fig. 3D)**. Surprisingly, we find that with this procedure, networks were unable to learn any of the target functions with the same efficiency as with resonant learning (**Fig. 3E, S19)**. Even for the most straightforward task of the three-input learning case, the resonant learning scheme converges at least one order of magnitude faster than learning with a non-oscillating hub. We attribute poor learning with a fixed hub state (or followin the dynamic rule in Eq.(1)) to the network’s inability to learn longer target cycles without an oscillating, periodic input that primes the network with a timescale that matches the length of the target time series.

Next, instead of using square wave function we ask whether other oscillatory patterns could give an advantage in learning. First, we test a simple learning case in which the networks must learn three different target functions of lengths {*L*_1_, *L*_2_, *L*_3_} = {8,10,12}. As shown above, networks can easily learn these different target functions when we use matching square wave input periods {*T*_1_, *T*_2_, *T*_3_} = {8,10,12}. By contrast, we show that when we use periodic random patterns of 0 and 1s of length {*T*_1_, *T*_2_, *T*_3_} = {8,10,12}, instead of square waves, the network can still learn efficiently, indicating that the timescale, not the specific repeated pattern, associated with the oscillations of the input may be the key feature for resonant learning (**Fig. 4A, B)**. However, an alternative explanation may be that the distinct input functions throughout each evolutionary run, unique to each target function could govern learning. To rule out this hypothesis, we then generate three distinct white-noise signals with no specific timescale used as input functions of the hub to learn three different target functions of lengths {*L*_1_, *L*_2_, *L*_3_} = {8,10,12}. We find that learning performance is degraded when we use white noise input functions in the absence of typical timescale (**Fig. 4A, B)**. Altogether, these results led us to propose that the well-defined timescale associated with the input period is the key feature that governs resonant learning. Because the specific pattern of the oscillatory input signal at the hub is not central for the network to learn the target function efficiently, we ask if the network is capable of learning multiple target functions of the same length as a function of different patterned inputs in one evolutionary run. Indeed, we find that the networks can learn multiple target functions to match inputs all of length *10* **(Fig. S18)**. By contrast, when the targets vary in length, the network fully learns the shorter length targets before learning the longer period target functions (**Fig. 3B-D**).

**Figure 4.**
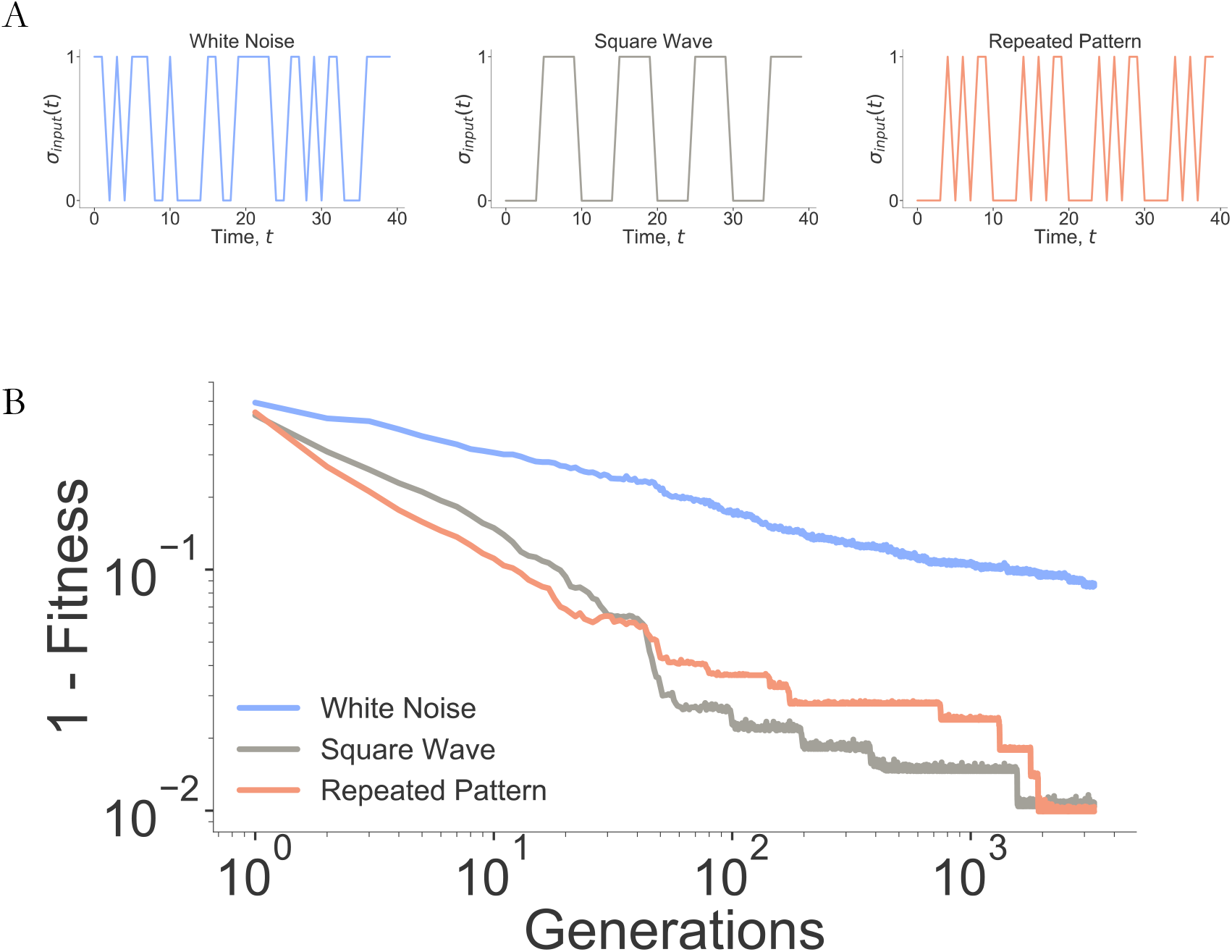
Learning is solely dependent on the timescale of oscillations. **(A) Different periodic hub input time series**. We define three different input types: Square wave inputs which we have used previously, a white noise pattern with a set sequence and no characteristic timescale, and a repeated pattern input with a fixed period. (**B) Period matters, not patterns**. We compare multiplexed learning of three different target functions of length 8, 10, 12 for networks *N* = 500, *γ* = 1.9 on the three types of input sequences. Repeated pattern inputs and square wave inputs yield faster learning than white noise inputs, showing that the timescale associated hub oscillations matter more than the specific repeated pattern. In these simulations.

## DISCUSSION

In this work, we demonstrate that by forcing the network hub to oscillate we can induce a wide variety of novel attractor cycles. This finding opens the door to new strategies about how to control complex network behaviors with access to only few nodes such as hubs. Furthermore, we show that this property can be used in a network-based learning procedure for which a single Boolean network can be trained to learn multiple targets. Surprisingly, *resonant learning* outperforms similar learning procedures in the absence of oscillating inputs for comparable tasks by orders of magnitude. *Resonant learning*, however, is not the only way for networks to learn multiple targets. Previous studies demonstrated that topological separation into distinct domains of the network architecture was necessary for learning multiple tasks (31, 32). It is therefore interesting to note that learning multiple task is facilitated by either behavior.

While *resonant learning* is a potentially useful property, in this work we have not directly applied it to known machine learning frameworks such as reservoir computing. However, it has already been shown that Boolean networks whose dynamics are tuned at the vicinity of the critical regime can produce more flexible reservoirs (4, 5), consequently, it would also be interesting to assess in the future how using oscillating hubs from reservoirs with a scale-free topology could affect the learning efficiency in such frameworks.

Finally, given the ability of oscillating hubs to govern network-level behavior through a frequency domain, we posit that frequency-based coordination might also be a mechanism in real biological systems as suggested by recent experiments (11) (12). More generally, our study sheds light on a novel property of large dynamical systems and how we may exploit it to improve learning of dynamical behaviors.

## ACKOWLEDGEMENTS

We thank Yaron Singer and Nimrod Shaham for their feedback on an earlier version of this work. Authors thank Mayra Garcia for early assistance. We also thank the members of the Cluzel lab for discussions and feedback. This work was funded in part by the Harvard Department of Systems Biology Summer Research Program and Harvard Microbial Sciences Initiative Summer Research Fellowship to SG. We are grateful for access to the Harvard Odyssey compute cluster on which these simulations were performed. M.A. acknowledges PAPIIT-IN111322 UNAM grant for partial support. P.C. acknowledges partial support from NSF grant 1615487.

## Author Contributions

S.G. and P.C. designed research; S.G. and M.A. performed research; S.G., M.A. and P.C. analyzed data; and S.G., M.A. and P.C. wrote the paper.

## Footnotes

a For clarity, we define resonant frequencies in the network as multiples of the input oscillation frequency,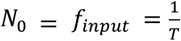. We can convert any resonant frequency back to the time domain by inverting it, giving resonant periods For period *T = 10*, the input frequency is *N*_0_ = 0.1, and a resonant frequency is 5*N*_0_ = 0.5, which corresponds to a resonant period *T*_*res*_ = 2.

