## Supplemental figures for "Resonant Learning in Scale-free Networks"

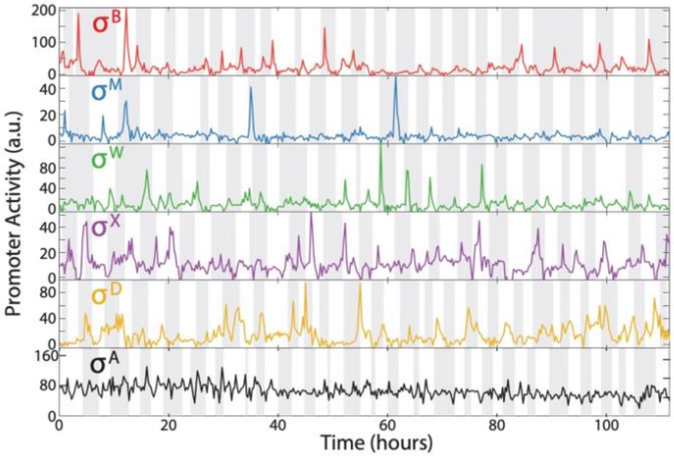


**Figure S1.** Promoter activity time series of central regulatory transcription factors in single cell gene expression experiments conducted in *Bacillus subtilis*. As shown, these transcription factors pulse over time, exhibiting an oscillatory behavior. Reproduced from Park et al. Cell Systems 2018.

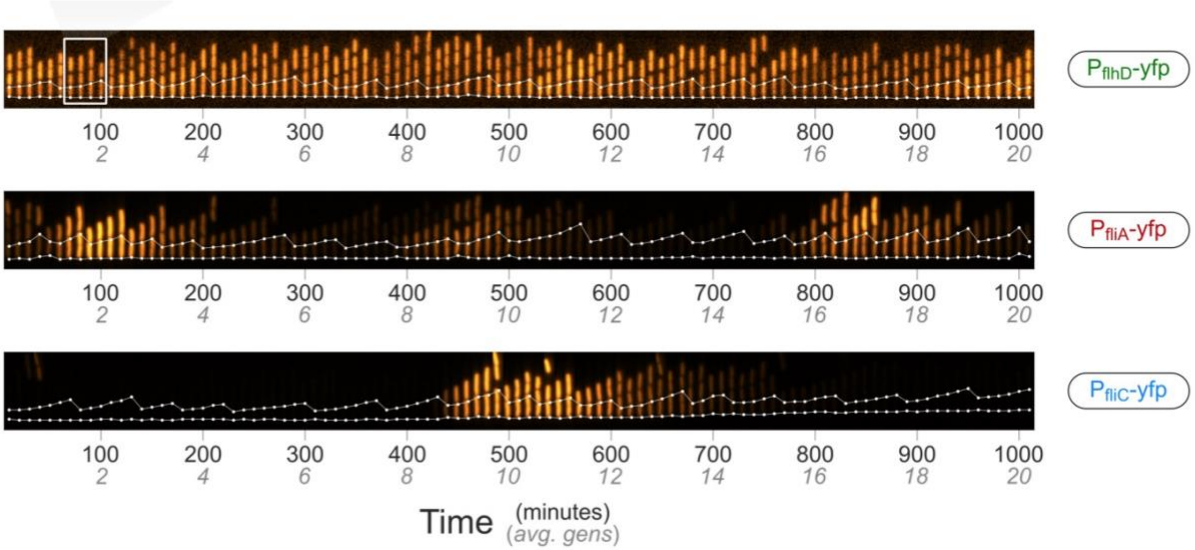


**Figure S2.** In a separate set of experiments conducted in *E.coli*, the promoter expression of flhD, fliA, and fliC, central genes responsible for flagella biosynthesis, also pulse with time at varied frequencies. This suggests evidence for pulsatile expression patterns in existing biological systems. Figure reproduced from Kim et al. Science Advances 2020.

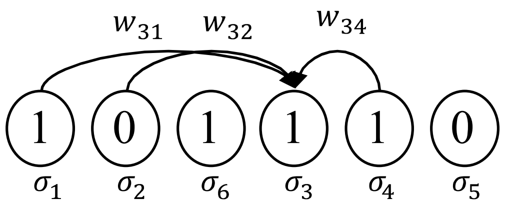


$$\sigma_i(t+1) = \begin{cases} 0, \sum_{j=1}^{N} w_{ij}\sigma_j < 0 \\ 1, \sum_{j=1}^{N} w_{ij}\sigma_j > 0 \\ \sigma_i(t), \sum_{j=1}^{N} w_{ij}\sigma_j = 0 \end{cases}$$

**Figure S3. Dynamical updating rule**. We define the network to have a simple thresholded activation function to determine the state of the node $\sigma_i(t+1)$ as defined in Eq.(1) of the main text. For each connection $\sigma_i \rightarrow \sigma_j$, the connection weight $w_{ij}$ is randomly chosen with uniform probability in the interval [-1,1] and kept fixed throught the temporal dynamics of the network. To determine the value $\sigma_i(t+1)$ for each node in the network at time $t+1$, we have to know the states at time $t$, $\sigma_j(t)$, and the weights of all nodes that have an out-going edge to $\sigma_i$. While simple, this updating rule has several advantages over the more classical Kauffman networks, which require a truth table of memory $O(N2^K)$ to define the updating rules for each of the $N$ genes in the network (each with average connectivity $K$), allowing us to simulate larger random networks efficiently.

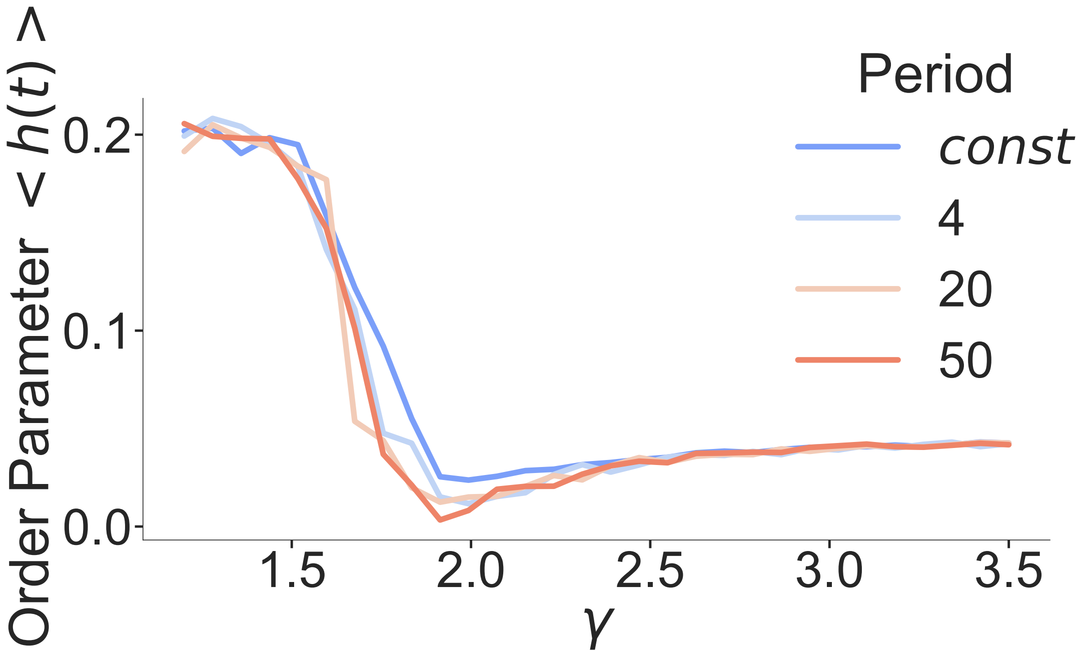


**Figure S4.** Threshold networks exhibit a continuous phase transition from ordered to chaotic states. As the scale free parameter $\gamma$ increases, the average connectivity of the network decreases and the network becomes less chaotic, transitioning to an ordered regime around $\gamma = 1.9$. Because the threshold activation function makes analytical approximation intractable, we instead evaluate this phase diagram empirically through simulations using the Hamming distance as the order parameter: $\lim_{t\to\infty} h(t) = \lim_{t\to\infty} \frac{1}{n} \sum_{i=1}^{n} |\sigma_i(t) - \tilde{\sigma}_i(t)|$. We perturb a fraction, $d$ = *0.05* of the states in the network and calculate the difference in trajectories between the perturbed initial condition, $\tilde{\boldsymbol{\sigma}}(\boldsymbol{t})$, and the original condition, $\boldsymbol{\sigma}(\boldsymbol{t})$. For each $\gamma$, we average $\langle h(t) \rangle$ over five different initial conditions and twenty different networks. Interestingly, this phase transition is preserved even when the hub of the network oscillates at a given frequency. As confirmed by Zañudo *et al.*, these Boolean threshold networks are such that $\langle h(t) \rangle$ approaches 0.2 in the chaotic regime, rather than 0.5, due to many nodes in the network freezing in either the 0 or 1 state. We calculate the phase transition diagram for these networks when the most connected node (the hub) in the network oscillates with different periods to confirm that this does not change the ordered-chaos phase transition.

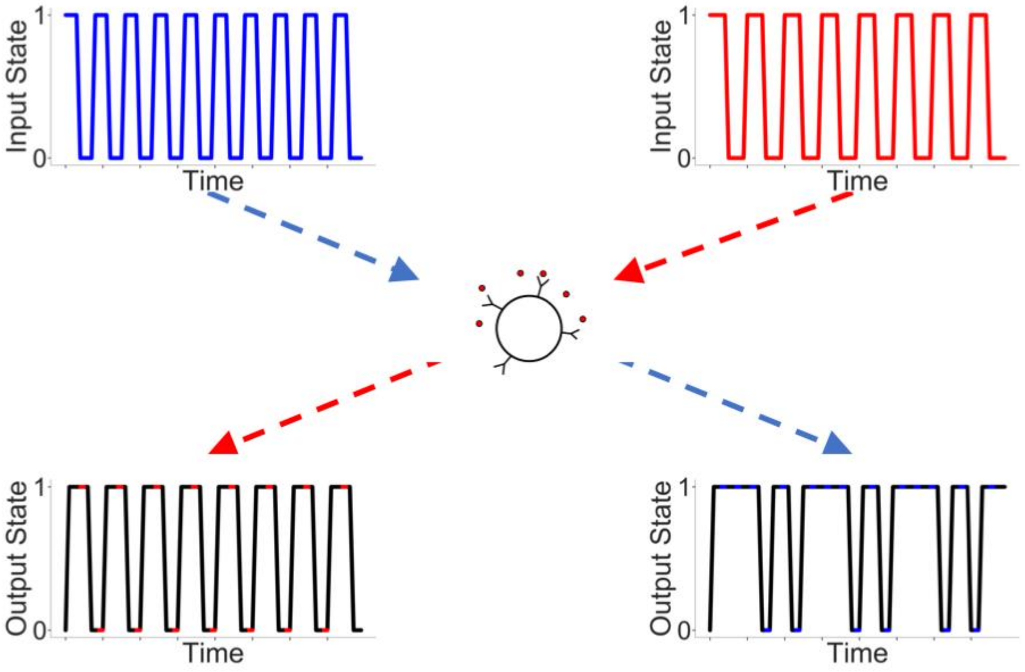


**Figure S5.** The corresponding time series for the frequency domains shown in Figure 1B. Here, we color the output states when they are synchronized with the input state.

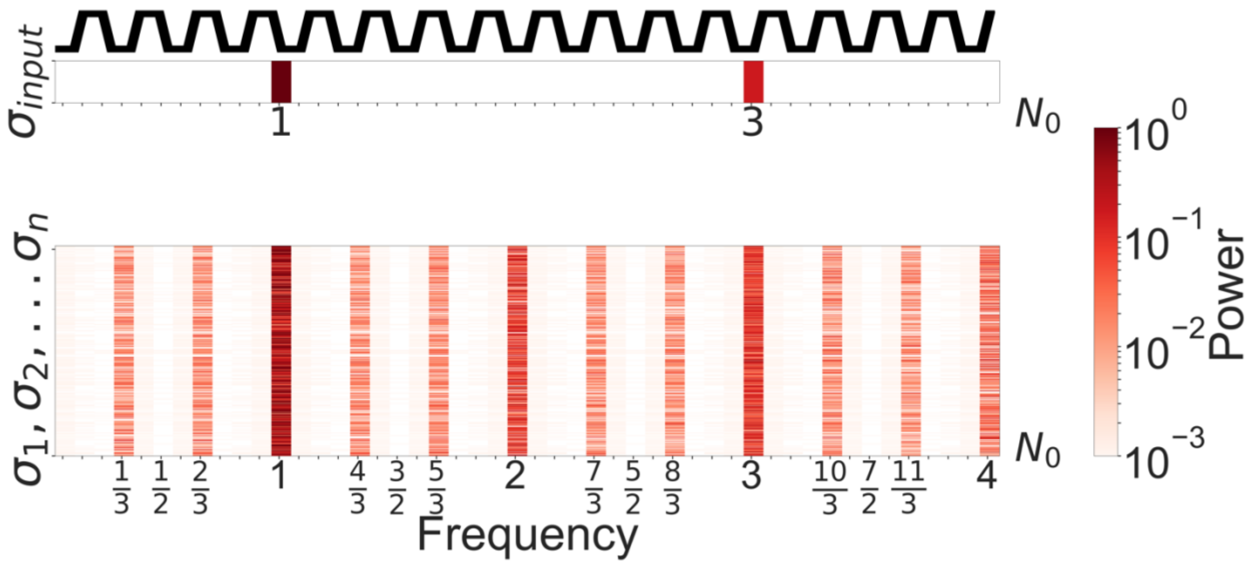


**Figure S6.** The power spectra of input and network nodes, as shown in Figure 1C, with an input period of T=8. To calculate the power spectra for a given node's time series in the Boolean networks, we sample only the parts of their time series in which the node is in a known attractor. We extract the states of a given node**,** $\sigma_i(t_{start}), \sigma_i(t_{start}+1), \ldots, \sigma_i(t_{end})$ , We use the built-in *Periodogram* function in Scipy's signal library of Python to estimate the power spectra of these signals for comparison. We plot only the dominant frequency.

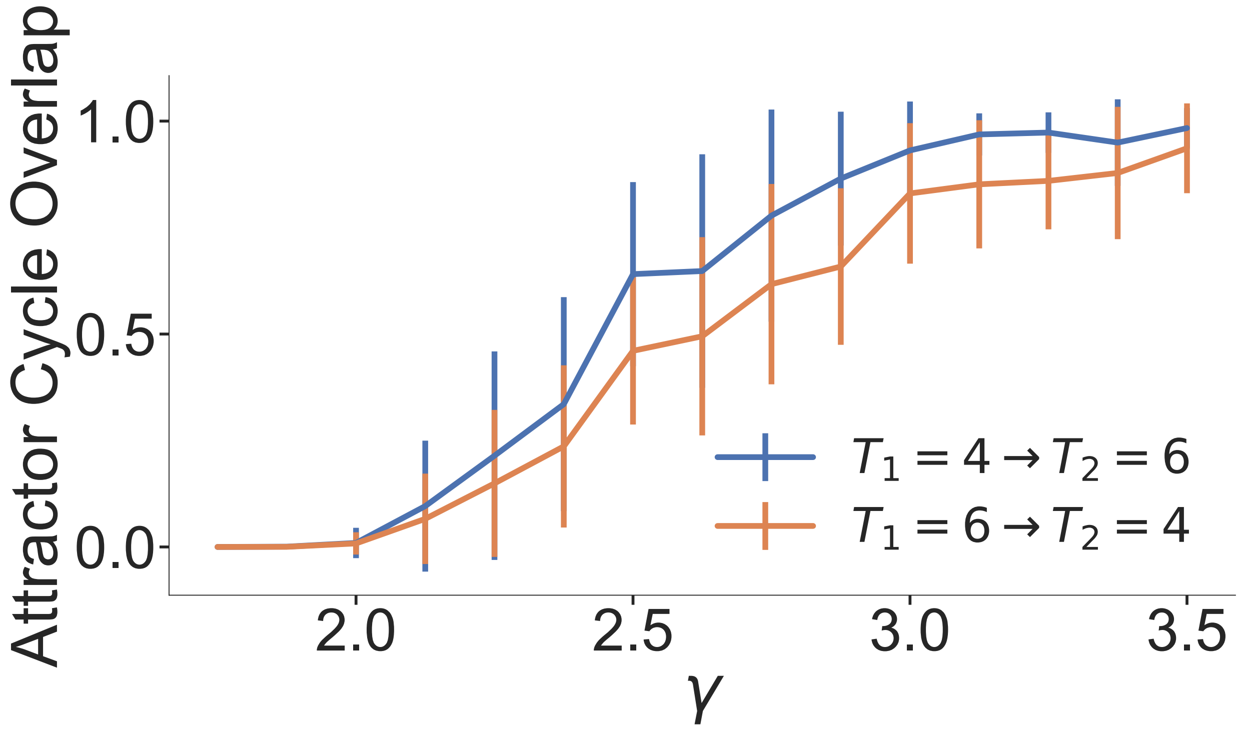


**Figure S7: Overlap between new attractors states.** When creating new attractor cycles with oscillations, we considered how different the newly created attractor cycle states were with respect to each other. It would be uninteresting if oscillating the input node at two separate frequencies resulted in two attractors incorporating the same network states. We tested this by generating 20 different networks for each gamma value and two oscillation periods, T=4 and T=6. We probed the attractor landscape to identify up to 20 attractors and, for each period, found the most similar attractor cycle in the landscape corresponding to the other period. We find that for $\gamma < 2.3$, the average overlap between attractors created by a slight difference in input oscillation is low, indicating that we generate novel attractor cycles for different input signals.

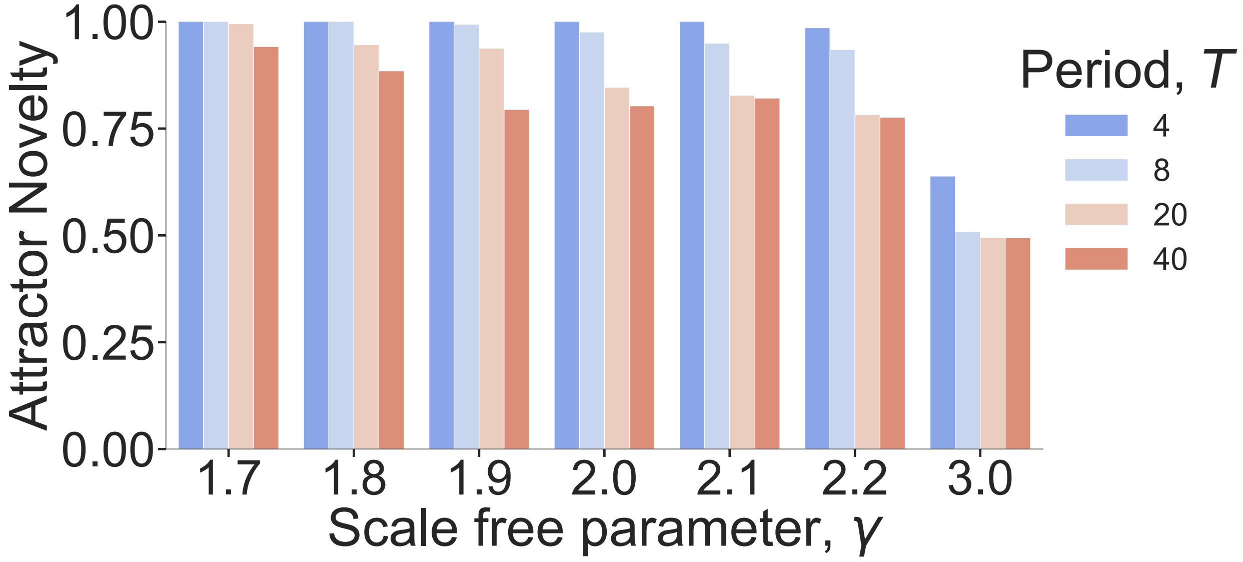


**Figure S8.** In addition to cataloging similarity between the new attractor states for different periods of input oscillation, we compare the new attractor states to attractor states when the input, $\sigma_{input}$ is blocked. Let $A_T$ be the set of all network configurations $\boldsymbol{\sigma_i}$ such that $\boldsymbol{\sigma_i}$ is part of an attractor cycle when $\sigma_{input}$ oscillates with period $T$, and let $B$ represent the set of all network configurations $\boldsymbol{\sigma_j}$ such that $\boldsymbol{\sigma_j}$ is part of an attractor cycle when $\sigma_{input}$ is blocked at OFF or ON. We calculate $1 - \frac{|A_T \cap B|}{|A_T|}$ averaged over 10 different networks and 1000 initial conditions for each value of $\gamma$ to identify the attractor landscape. As the input period increases beyond the relaxation timescale of the network, there is always some overlap with the control landscape.

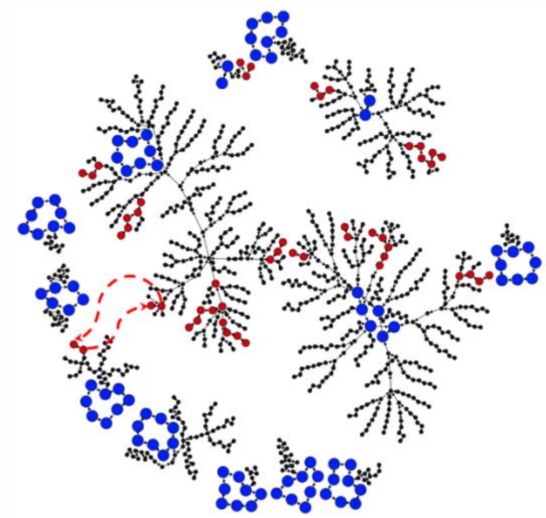

**Figure S9: Visualizing the attractor landscape.** For a sample network with $N = 1000$ and $\gamma = 1.9$, we probed the network's attractor landscape in the conditions where the input node is set ON and then OFF. Each circle represents a network state, and an arrow indicates a transition based upon the updating rule from one state to another. The attractor cycles are shown in blue for this network when the input is blocked at either an ON or OFF state. We oscillated the input at varied periods between the ON and OFF states and found new attractor cycles, whose corresponding states are colored in red. We highlight one particular new attractor cycle of an input oscillation of T=4. **Phase robustness.** We note that, unlike normal, fully deterministic Boolean threshold networks, a given network state in these networks can be part of multiple basins of attraction in our new model. That is, depending on the phase of the input node, or what part of its switching behavior it is in, the network state can converge on different cycles.

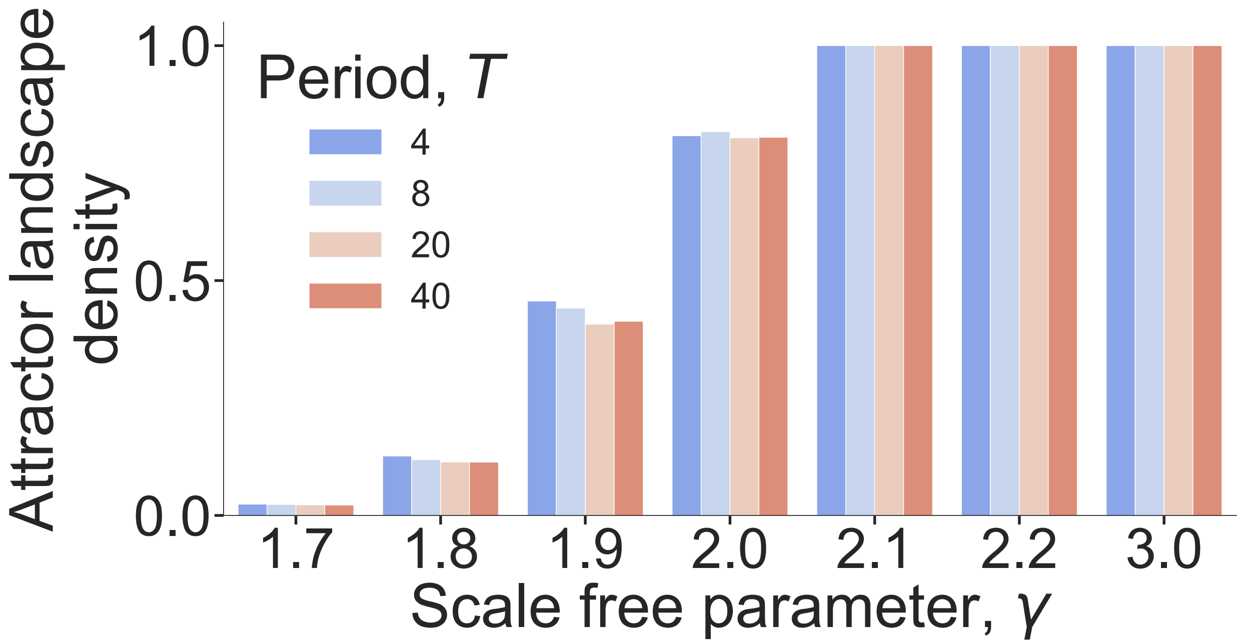


**Figure S10.** New attractor cycles are robust**.** We quantify in 1,000 initial conditions how many unique attractor cycles are found. Again, we average over 10 different networks for each $\gamma$. We find that the new attractor cycles constructed from input oscillations in low-$\gamma$ networks are indeed robust, as the attractor landscape is sparse. Note, for clarity of presentation, we add 0.02 to the bars for gamma=1.7 in order to see that these values are non-null. There are indeed some attractors discovered.

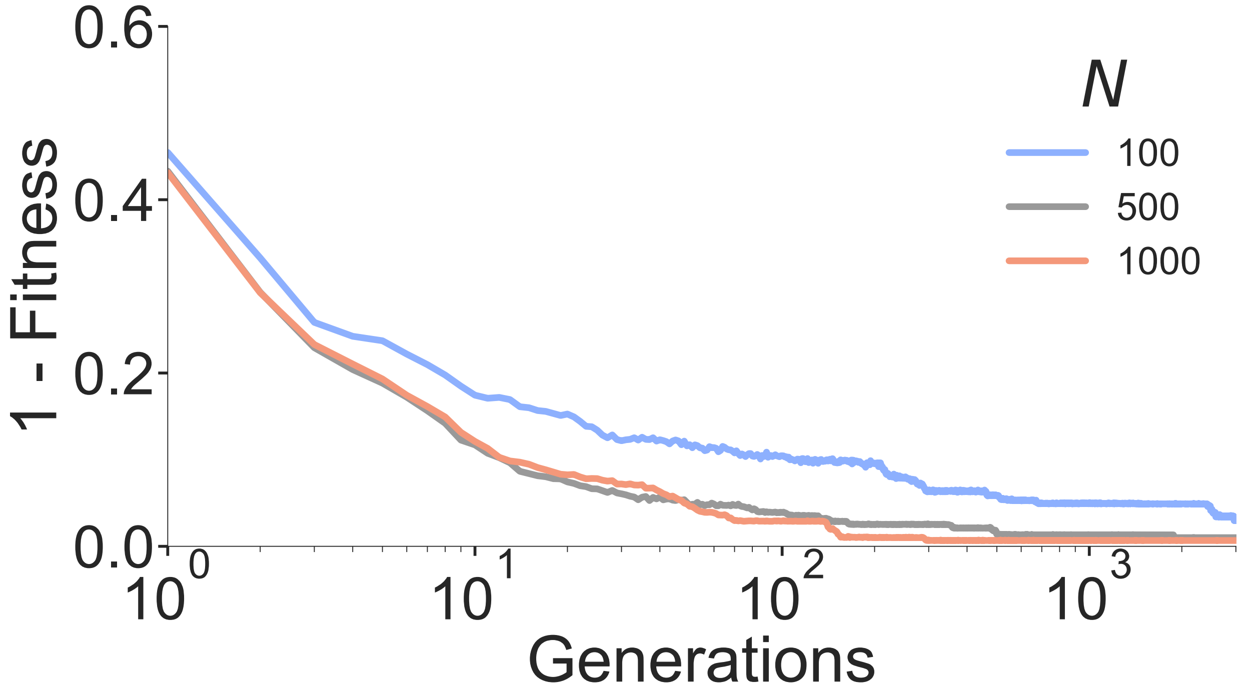


**Figure S11.** Error in the learning process of the network throughout generations (see also Fig. S14). The learning consists in training the network so that the temporal signal of one node chossen randomly, (the output node), matches a predefined function (the target function). The error ($E = 1 - Fitness$) is the difference between the actual temporal signal of the output node and the target function (see Fig. S14). It can be obseerved that thoughout generations the error decreases, which means that the networks learn to reproduce the target function with increasing precision. We report here the results for networks of different size. In each case the network was trained to pair three different input signals of period $T = \{6,8,10\}$ to match targets target functions with the same periods. This results demonstrate that networks with $N = 500$ and $N = 1000$ learn essentially at the same rate.

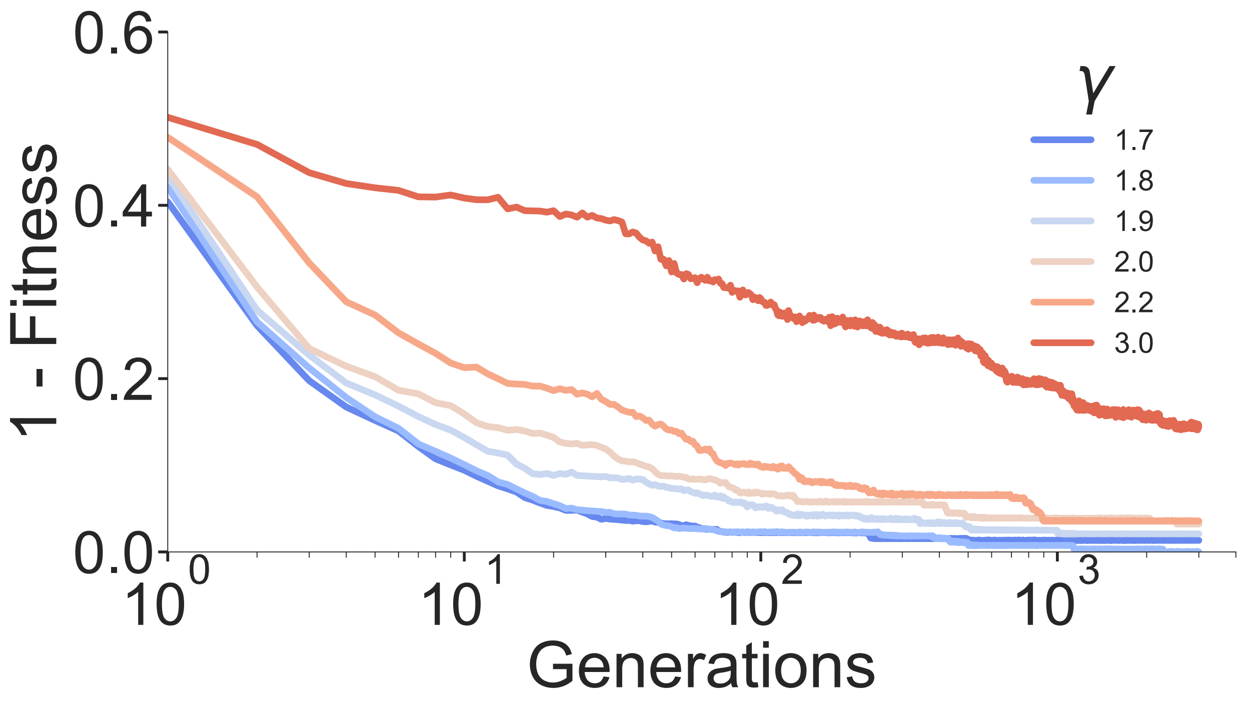


**Figure S12: Testing multiple gamma values.** Error in the learning process throughout generations for networks with different values of the scale-free exponent $\gamma$. The learning process is the same as described in the main text and in Figs. S11 and S14. It can be observed that networks with lower values of $\gamma$ are able to learn significantly faster, (particularly for $\gamma < 1.9$), than networks with larger vaues of $\gamma$, justifying our choice of $\gamma = 1.9$ in most of our simulations.

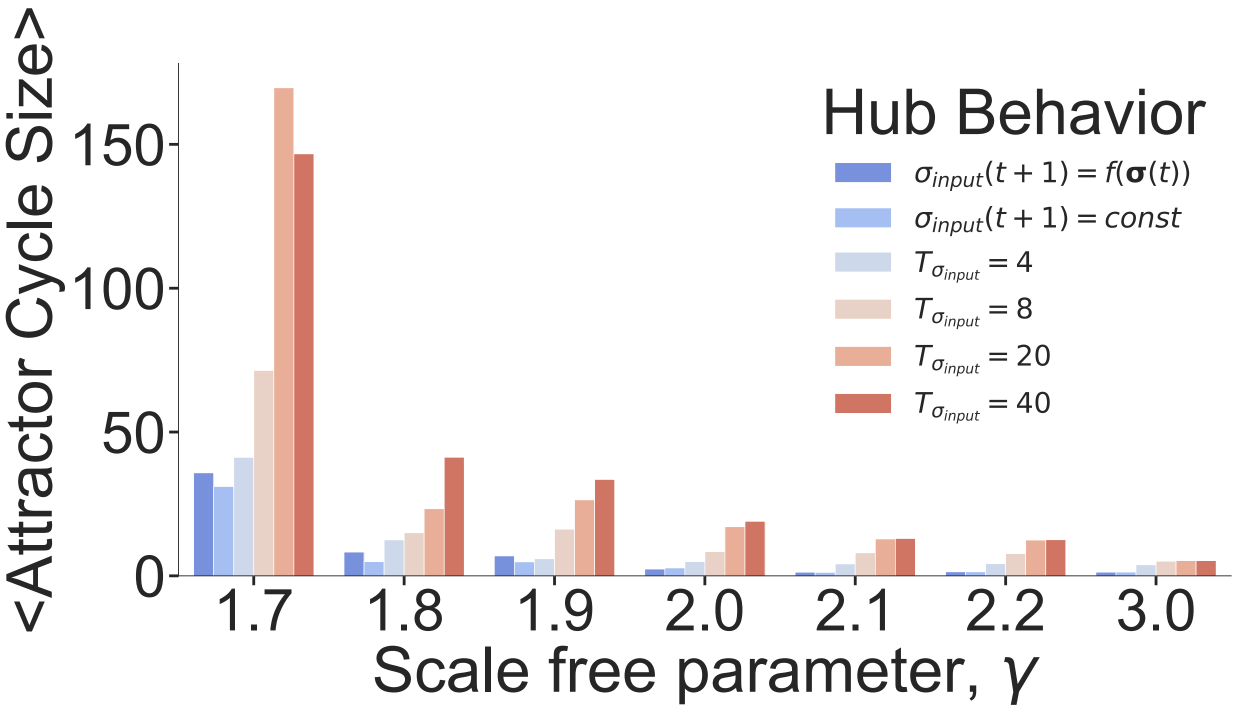


**Figure S13: Average size of attractor cycles**. We catalogue in the probed attractor landscapes the average size of each attractor found. In chaotic networks, the size of the attractors is far larger than in ordered networks, becoming unreasonable to simulate for $\gamma < 1.7$. Additionally, we show the average attractor size when the hub node is allowed to update naturally as a function of other nodes in the network and when the hub input node is set to a constant, blocked value of ON (OFF).

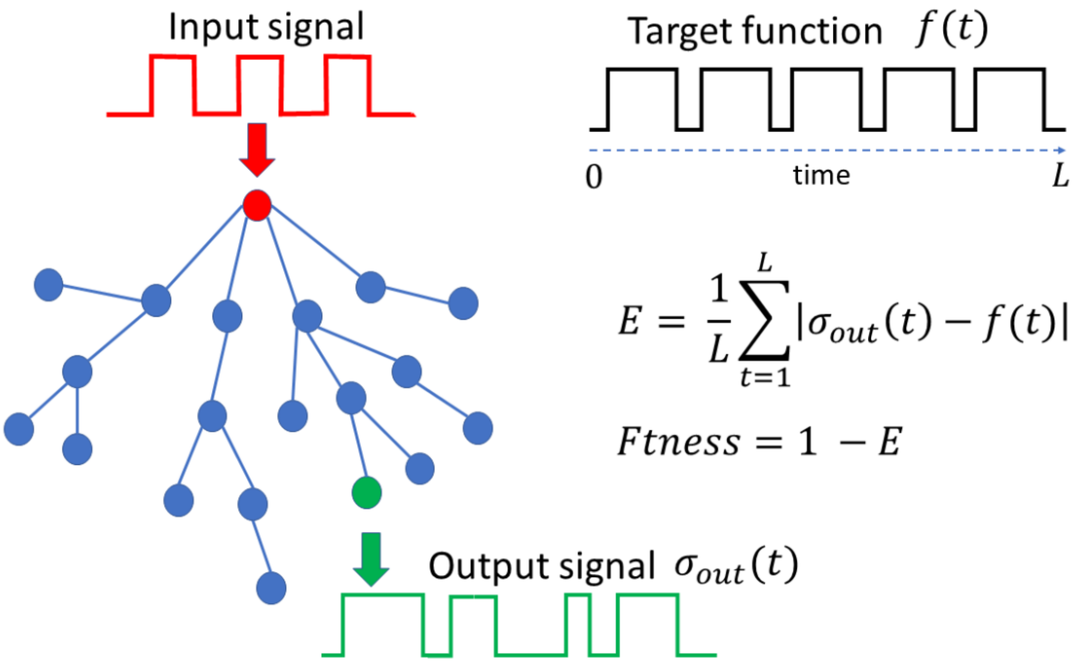


**Figure 14. Schematic illustration of the network learning process.** A periodic input signal with period $T$ is introduced into the hub of the network (red node). A randomly selected node (other than the hube) is chosen to be the output node (green node). During the temporal dynamics of the network (Eq.(1) of the main text) the output node will generate an output signal $\sigma_{out}(t)$ that depends on the specific details of the networks (connections and weights), and on the form of the input function. The output signal $\sigma_{out}(t)$ is then compared with a predefined target function $f(t)$ which can or cannot be periodic (in the illustration, the target function is periodic, but this is not necessaryly the case). Both the input and target functions are computed for a period $L$ of time. The error $E$ between $\sigma_{out}(t)$ and $f(t)$ is computed as the normalized Hamming distance between these two functions averaged over the time interval $L$.

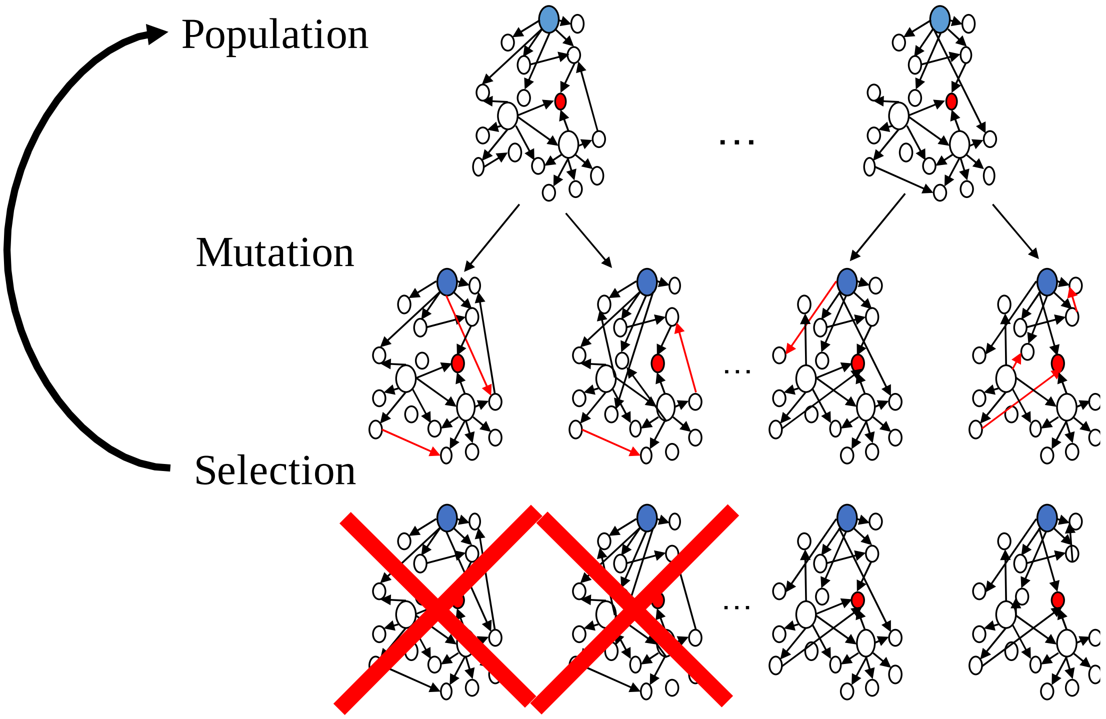


**Figure 15. Schematic illustration of the evolutionary algorithm.** Beginning with a population of $N_{pop} = 50$ networks, we generate three mutants for each network. Murations consist in randomly rewiring the network conections and/or changing the connections weights. The mutation rate is $\mu = 0.02$ for each node in the network. Then, we calculate the fitness function of each of our networks in the population, and select only the best $N_{pop} = 50$ networks, which are the ones that pass to the next generation. This reduces the population to its original size. One full process of mutations and selection represents one generation.

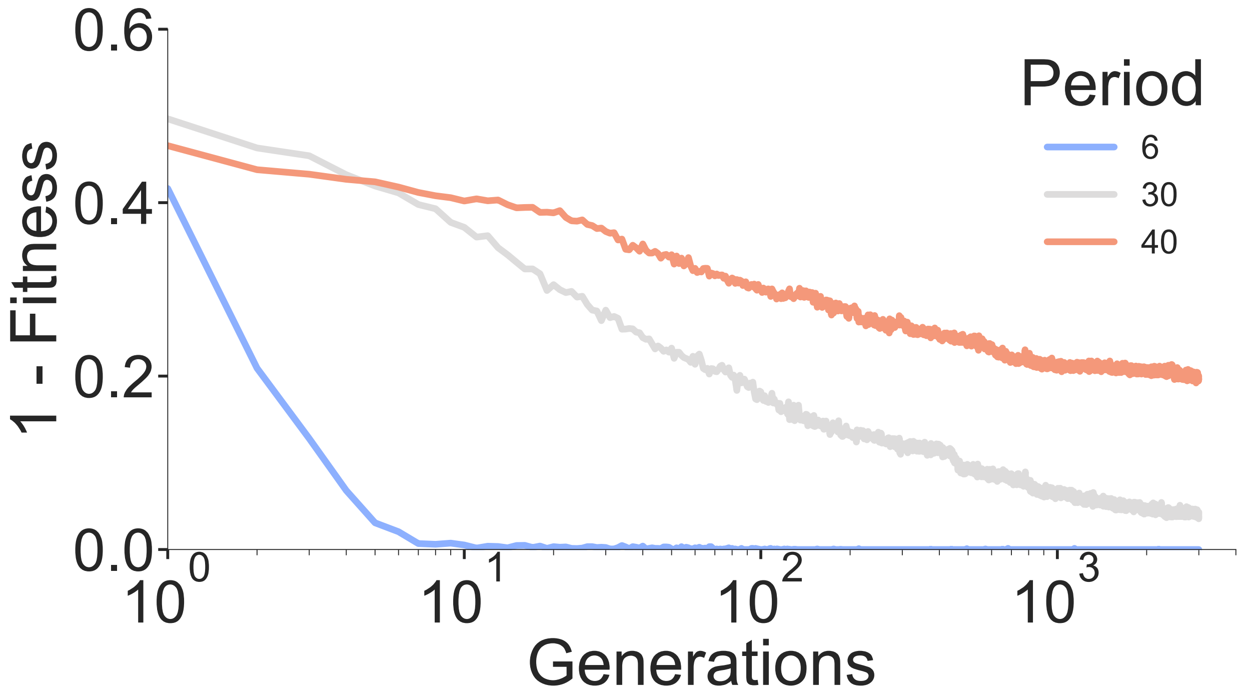


**Figure S16. Limitations on learning.** We demonstrate that networks with gamma N = 500, $\gamma$ = 1.9 struggle to learn specific attractor cycle behaviors for long time scale input periods. Specifically, we assayed learning for input periods, *T*, and target cycle length, *L*, *T*=*L*={6,30,40}. As can be seen, the network is only able to completely learn the short timescale of *T* = *L* = 6, demonstrating the limitation of resonant learning for targets longer than the relaxation time ($t_{relax}$=20) of the network.

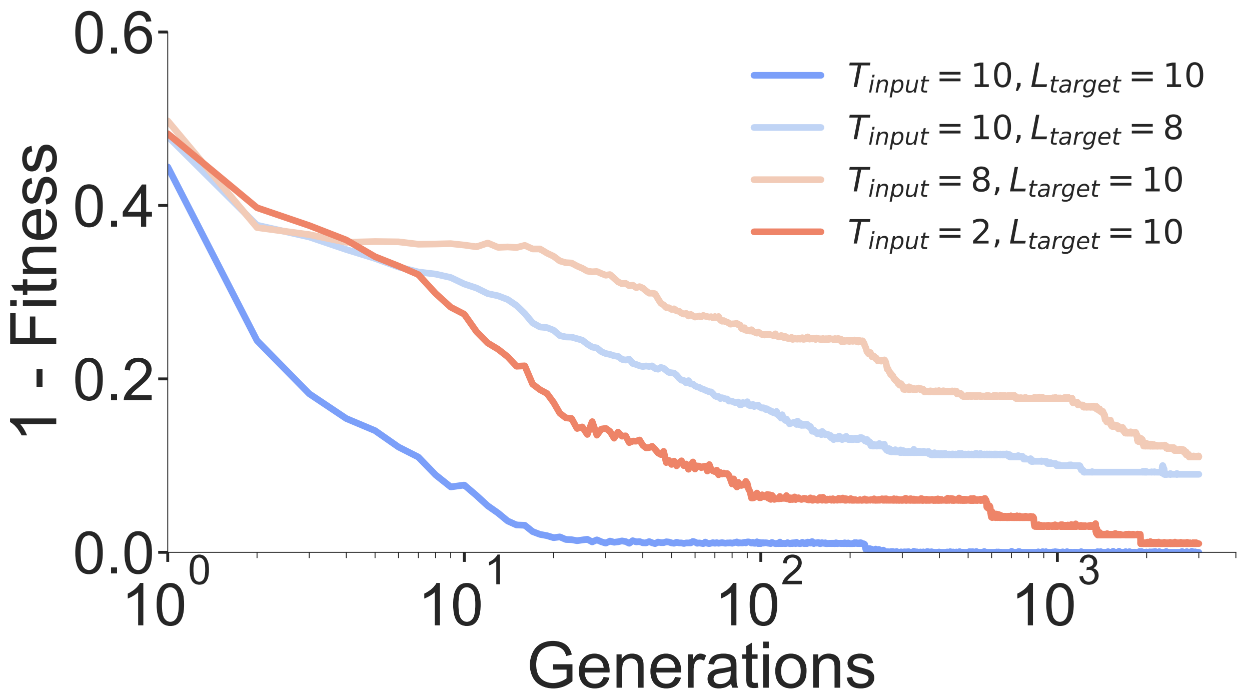


**Figure S17.** Learning target functions with lengths that differ from the period of the input function. Here, we find that the network learns best when the input period and target are of equal duration. Interestingly, even having a divisor of the target cycle length does not yield the same benefit in learning, as is the case for $T = 2, L = 10$). Learning performance worsens when the target is longer than the input period and is not a multiple of it ( $T = 8, L = 10$). We have a slight improvement in learning performance when the target period is shorter than the input length ($T = 10, L = 8$).

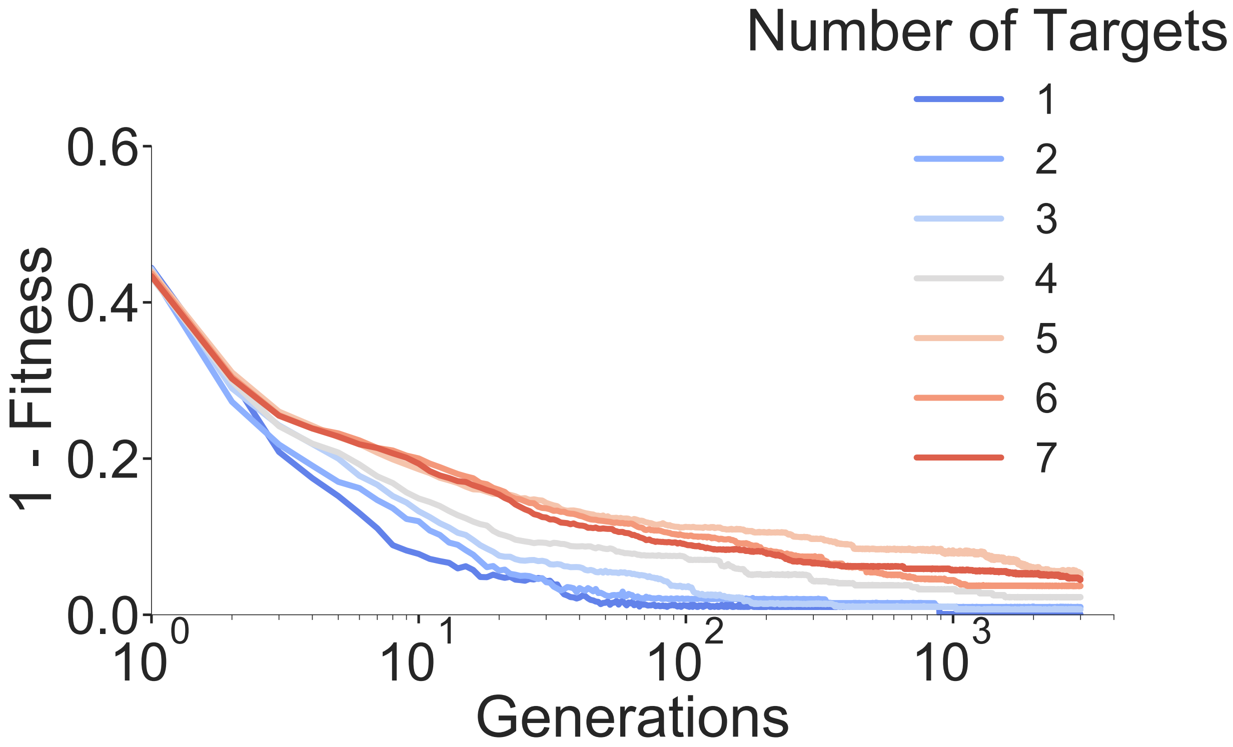


**Figure 18. Learning several patterns.** We force the network to learn repeated input functions of size $T = 10$ to output functions of the same length $L = 10$. Because the inputs are arbitrary repeated patterns, our fitness function can allow the network to learn different repeated patterns of the same length without compromising other repeated pattern inputs. This result shows that the network's ability to learn an arbitrary number of repeated patterns considerably degrades when the number of patterns is larger than 7.

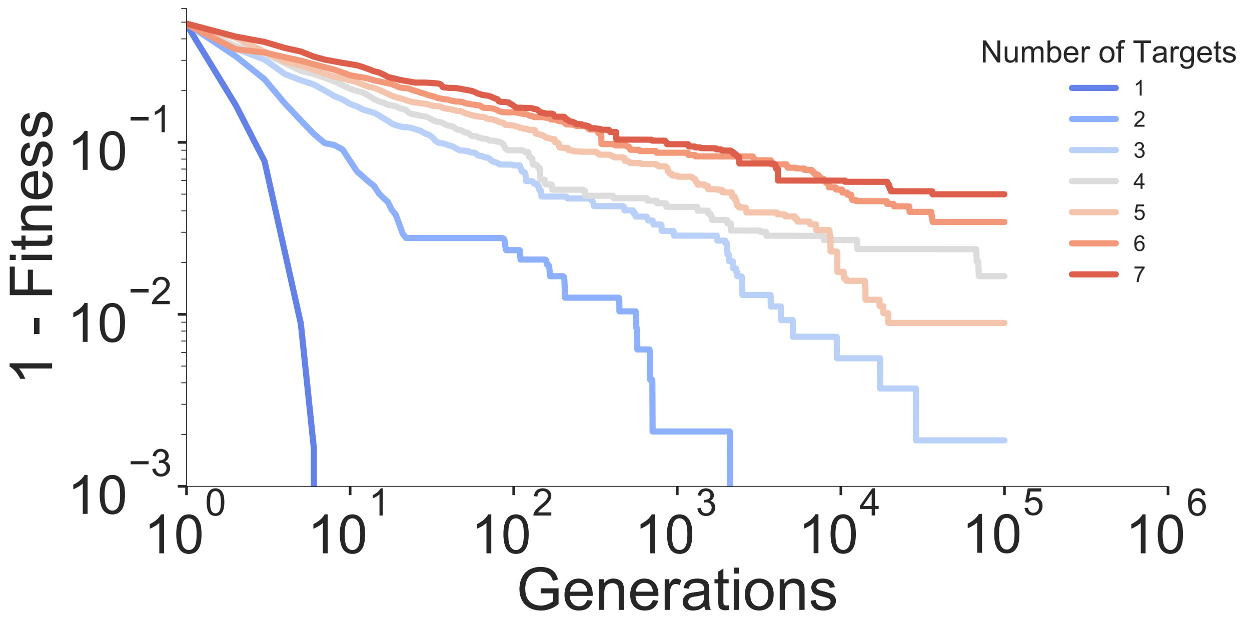


**Figure S19. Learning multiple target functions of varied length when the input node is under the same dynamical rule as the rest of the network**. We repeat the same procedure as for Figure 3D but allow the input node's dynamics to follow the same updating rule as the rest of the network (Eq.(1) of the main text), giving the network more control over its behavior. We seek to level the playing field between the oscillating input case, where the network has a signal from the hub (Figs. 3C and 3D of the main text). While learning improves, the network cannot learn long targets, yielding poor performance when learning multiple attractor functions.

**Defining an attractor.** Since both the network connections $\sigma_i \rightarrow \sigma_i$ and the connection weights $w_{ij}$ are determined when the network is constructed and do not change in time, the network dynamics given by Eq. (1) of the main text are deterministic. Additionally, a network with $N$ nodes has a finite number of $2^N$ states. The configuration space is finite. Therefore, after some transient time, the network inevitably will fall in a previously visited state. The dynamics will repeat from that point on, making the network fall into a periodic pattern of activity. In the first part of our results, when discussing the attractors of the network, we include the input node as part of this attractor definition. Therefore, if the input node oscillates with an input period $T = 4$, the attractor must, by definition, be a multiple of 4. To allow for more general results, when forcing these networks to learn different attractor states, we remove this restriction from the definition of the attractor. We generally define the attractor as the expression pattern of all nodes, excluding the input node (the hub).
